# Task-irrelevant features in working memory alter current visual processing

**DOI:** 10.1101/2024.10.29.620777

**Authors:** Hongqiao Shi, Qiqi Zhang, Jieyudong Zhou, Yue Ding, Yonghui Wang, Ya Li

**Author notes:** authors contributed equally (Hongqiao Shi, Qiqi Zhang). Yonghui Wang, co-corresponding author. Ya Li, corresponding author.

## Abstract

Higher-level cognition depends on visual working memory (VWM), the ability of our brain to maintain and manipulate internal representations of images that are no longer presented to us. An important question in this field is whether VWM is represented in a sensory or nonsensory manner. Progress has been made in understanding the features to be remembered, but the representational nature of the memory-irrelevant features is unclear. Here, we used a dual-task paradigm to investigate how and when the memory-irrelevant features interact with the concurrent visual information. In a series of experiments, participants were asked to perform a visual search task (Experiment 1) or a perceptual discrimination task (Experiments 2 and 3) involving a memory- irrelevant feature while simultaneously holding the other feature for later retrieval. Experiment 1 showed that VWM biases the allocation of attention to color matching to the memory-irrelevant color. More importantly, the degree of VWM-biased attention decreased monotonically with decreasing feature similarity, and this behavioral monotonic gradient profile resembled the tuning curve of feature-selective neurons in the early visual cortex. Experiment 2 revealed that irrelevant features biased ongoing perception, as indicated by the shifted discrimination threshold. Experiment 3 further demonstrated that VWM-biased perception occurs only at short delays but not at prolonged delays. Our results suggest that the memory-irrelevant feature is represented as a sensory analog for a limited period of time in the visual areas where it was initially processed. Our results extend sensory recruitment theory to memory-irrelevant features in VWM.

## Introduction

Working memory (WM), our ability to temporarily hold limited information and manipulate it in mind for future behavior, serves as a foundation for higher-level cognition (Baddeley & Hitch, 1974; Luck & Vogel, 1997; Mongillo et al., 2008). For example, when we try to find a green cup in the cluttered kitchen, the appearance of the cup (e.g., color and shape) could be maintained in visual working memory while concurrently looking at all the objects in the kitchen to search for the target. A fundamental question in the WM literature is how the brain represents objects that are not currently seen. Although multiple levels of cortical areas have been shown to be involved in the representation of VWM content (Christophel et al., 2017; LaBar et al., 1999; Miller et al., 1996; Xu, 2017; Yu & Shim, 2017), sensory cortices have been a concern over the past decade, which suggests the sensory analogy representation of VWM (Harrison & Tong, 2009; Huang et al., 2024; Pasternak & Greenlee, 2005; Rademaker et al., 2019; Serences et al., 2009). Accordingly, the sensory recruitment theory has been proposed, suggesting that VWM is maintained by the sensory cortical areas where VWM is initially processed. Behaviorally, previous studies have investigated the interaction between VWM and perception via a variety of perceptual tasks (Fang et al., 2019; Olivers et al., 2006; Pan et al., 2012; Soto et al., 2005; Teng & Kravitz, 2019). These behavioral studies further imply that the representation of WM in visual areas is behaviorally meaningful and supports the sensory analog representation of VWM. However, these studies have focused mainly on the memory-relevant feature, that is, the feature to be remembered by the object. The question is whether the memory-irrelevant features of the maintained object (hereafter referred to as irrelevant features) are represented as sensory analogs, which could complement VWM theories.

Information in VWM can influence the allocation of attention to the perceived stimuli in the environment, even when these visual stimuli are completely irrelevant to the task (Awh & Jonides, 2001; Olivers et al., 2006; Soto et al., 2005). It has been shown that irrelevant features could also bias attentional allocation, which suggests that irrelevant features of the maintained object could also be represented (Foerster, 2017, 2020; Foerster & Schneider, 2019; Gao et al., 2016; Kerzel & Huynh Cong, 2021; Olivers, 2011; Soto & Humphreys, 2009). However, VWM-biased attention itself cannot indicate sensory or nonsensory representations. Studies of visual attention have revealed monotonic gradient profiles (Mangun & Hillyard, 1988; Wang et al., 2022) or center- surround profiles of the attention effect (Hopf et al., 2006; Störmer & Alvarez, 2014) to prioritize the attended target from distractors. Intriguingly, VWM faces the same situation that demarcates the remembered stimuli from perceived stimuli; an important question in this regard is whether and how these profiles are at play within VWM. This issue is particularly important since such a profile in feature space could offer us a unique opportunity to gain insight into the representational nature of VWM(Fang & Liu, 2019; Kiyonaga & Egner, 2016; Teng & Kravitz, 2019). Neurons in the early visual cortex respond selectively to their preferred feature and monotonically decrease in firing rate as they move further away from the preferred value in the feature space (Bohon et al., 2016; Hubel & Wiesel, 1962; Johnson et al., 2008; Shapley et al., 2003). These feature-tuning properties have been attributed exclusively to the early visual cortex. If WM representations are sensory in nature, VWM-biased attention should reveal a monotonic gradient profile to prioritize the visual information that matches the VWM from others. Thus, the sensory analog representation account can be tested by assessing the profile of irrelevant feature-biased attention.

The VWM content has been shown to interact with the perception of the currently presented stimuli, which is in accordance with the behavioral prediction of the sensory representation hypothesis for memory-relevant features (Geng et al., 2017; Kang et al., 2011; Pan, 2020; Teng & Kravitz, 2019). For memory-irrelevant features, previous studies have provided evidence that irrelevant features can be maintained in the brain (Bocincova & Johnson, 2019; Zokaei et al., 2019). For example, Zokaei and colleagues have shown that the brightness of the maintained grating modulates the pupil response, even when the VWM task involves orientation recall and when the brightness of the retained object is a task-irrelevant feature. Additionally, an EEG study revealed the above-chance classification of EEG signals for irrelevant features in the early time window (Bocincova & Johnson, 2019). However, how memory-irrelevant features are represented in the brain has not yet been confirmed. Prominent theoretical accounts of VWM have proposed that multifeature objects are stored as integrated features (Luck & Vogel, 1997). If this holds true, the irrelevant feature might also be represented as a sensory analog and interact with ongoing perception. However, other evidence suggests that objects are maintained as independent individual features (Bays et al., 2011), which allows the sensory representation of irrelevant features to be verified. The sensory representation account was verified by examining whether the irrelevant features interact with real-time visual processing. If the irrelevant features in VWM are represented as sensory analogs and share a neuron population with visual processing, then the VWM representation would interact with the visual representation of the perceived stimuli, which in turn influences incoming perception.

To this end, we investigated the sensory analog representation account with two predictions addressed above. Experiment 1 examined the profile of VWM-biased attention in feature space. In particular, a search task was interleaved with the VWM maintenance period, during which the orientation of the multifeature object had to be recalled later. Importantly, we systematically varied the degree of similarity between the memory-irrelevant features (i.e., color) and the background context of the search targets (Experiment 1A) or distractors (Experiment 1B). We showed that VWM-guided attention decreased monotonically as the similarity between the memory-irrelevant feature and the perceived context feature decreased, which was consistent with the tuning curve properties of neurons in the early visual cortex. This raises the possibility that the same neural population in the visual cortex is recruited to maintain remembered memory-irrelevant information and to perceive incoming visual information. If so, we should observe the direct influence of the VWM information on the incoming perception. We tested this behavioral prediction of sensory representation in Experiments 2 and 3. During the maintenance period, a perceptual discrimination task was interposed. Importantly, we manipulated the similarity between the memory-irrelevant feature and the discriminative feature to examine the influence of VWM on ongoing perception. The dissimilar condition served as the baseline condition to measure the baseline discrimination threshold. If irrelevant features influence the incoming perception, then we should observe the shifted discrimination threshold in a similar condition relative to the dissimilar condition. Experiment 2 showed that the memory-irrelevant color (Experiment 2A) or orientation (Experiment 2B) shifted the interleaved discrimination threshold toward the irrelevant feature retained in WM, even when the color or orientation features were merely incidental and not required for the VWM task. Experiment 3 further showed that the VWM-biased perception induced by irrelevant features occurs at a 1 s delay but not at a 3 s delay, supporting the sensory representation for irrelevant features in VWM.

## Experiment 1: VWM-biased perception in the visual search task

Experiment 1 investigated whether and how feature similarity modulates the amount of attention biased by memory-irrelevant features. We had participants remember one feature (i.e., orientation) of the memory sample and perform a search task during WM retention. During the search task, the memory-irrelevant feature (i.e., color) was presented as the background of the search target (Experiment 1A) or distractor (Experiment 1B). If VWM content involuntarily influences concurrent visual processing, we would predict that the attention would be biased toward the color that matches or is similar to the memory-irrelevant feature, regardless of whether the color is the background of a target or a distractor, resulting in increased and decreased search speeds for target matching and distractor matching, respectively, in Experiments 1A and 1B. More importantly, we systematically varied the feature similarity between the maintained color and the perceived color. We reasoned that if the WM context is represented as a sensory analog, the degree of VWM-biased attention should decrease monotonically as the feature similarity between the maintained color and the perceived color decreases.

## Methods

### Participants

To determine the sample size and obtain adequate power, we used the power analysis tool G*power (Faul et al., 2007). To achieve 90% power with a significance level of 𝛼 = 0.05 and an effect size of partial eta squared 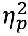= 0.06, the required minimum sample size was 26 for each experiment. The effect size was determined according to similar work on VWM-biased perception for relevant features(Teng & Kravitz, 2019). To ensure a sufficient sample size after data filtering, we recruited slightly more subjects than the calculated number. Experiments 1A and 1B recruited twenty-eight (20 females and 8 males; age: *M* = 20.0 years, *SD* = 2.3) and thirty participants (19 females and 11 males; age: M = 20.0 years, SD = 2.2), respectively. One participant was excluded because of overall low search accuracy, and two participants were excluded because of the experimenter’s maloperation, resulting in 27 subjects in Experiment 1B.

All participants had normal or corrected-to-normal vision, reported no history of mental illness, and were monetarily compensated for their participation. The study protocols were approved by the Ethics Committee on Human Experimentation of Shaanxi Normal University following the Declaration of Helsinki. Written informed consent was obtained from the participants before the experiments.

### Apparatus and stimuli

The participants viewed the stimuli in a dim room displayed on a 27-inch monitor with 2560 × 1440 resolution and a 100 Hz refresh rate. All visual stimuli were generated in MATLAB (MathWorks, Natick, USA) and Psychtoolbox-3 (Brainard, 1997; Kleiner et al., 2007). The participants viewed the stimuli at a distance of 70 cm, and a chin-and-head rest was used to stabilize their heads. All the stimuli were presented on a light gray background (grayscale = 128, luminance = 11.6 cd/m²). In Experiment 1, the memory sample was a colored grating (sample gratings: 2.5° in radius; spatial frequency: 2 cycles/degree) that contained two critical features: color and orientation. The color of the sample in each trial was randomly selected from a set of 40 colors (9° steps through the full 360° color wheel) evenly sampled in the CIE L*a*b color space. The color space was centered at L = 70, a = 0, b = 0, with a radius of 42 units. The orientation was assigned to the to-be-remembered feature and randomly chosen from 4 ranges (15° to 40°, 50° to 75°, 105° to 130°, and 140° to 165°) to avoid special orientations (e.g., vertical or horizontal). In the search display, two white-outlined squares (1.5° × 1.5°) with a 0.9° gap on one of four sides are presented on two colored circles (1.8° in radius) at an eccentricity of 8°.

### Procedure

The experimental procedure of Experiments 1A and 1B is shown in Fig. 1A. Each trial began with a 500 ms presentation of a fixation cross, followed by a 500 ms presentation of a memory sample. The participants were instructed to hold the orientation feature (i.e., relevant feature) of the memory sample for later reporting. The fixation cross was then presented for a 1000 ms delay, after which two search items were presented simultaneously until a response was made or 2000 ms had elapsed. The participants were asked to identify the target with the gap at the top or bottom by pressing "up" for the top gap and "down" for the bottom gap and were instructed to respond as quickly and accurately as possible. After a 500 ms interval, a vertical bar was presented at the center of the screen until the memory report or 3000 ms had elapsed. In this phase, participants could rotate the vertical bar clockwise or counterclockwise to report the remembered orientation as accurately as possible. The participants were instructed not to use gestures or any other strategies to complete the memory task, and they were supervised by the experimenter throughout the experiment. No feedback was provided in either the memory or the search task.

**Fig. 1.**
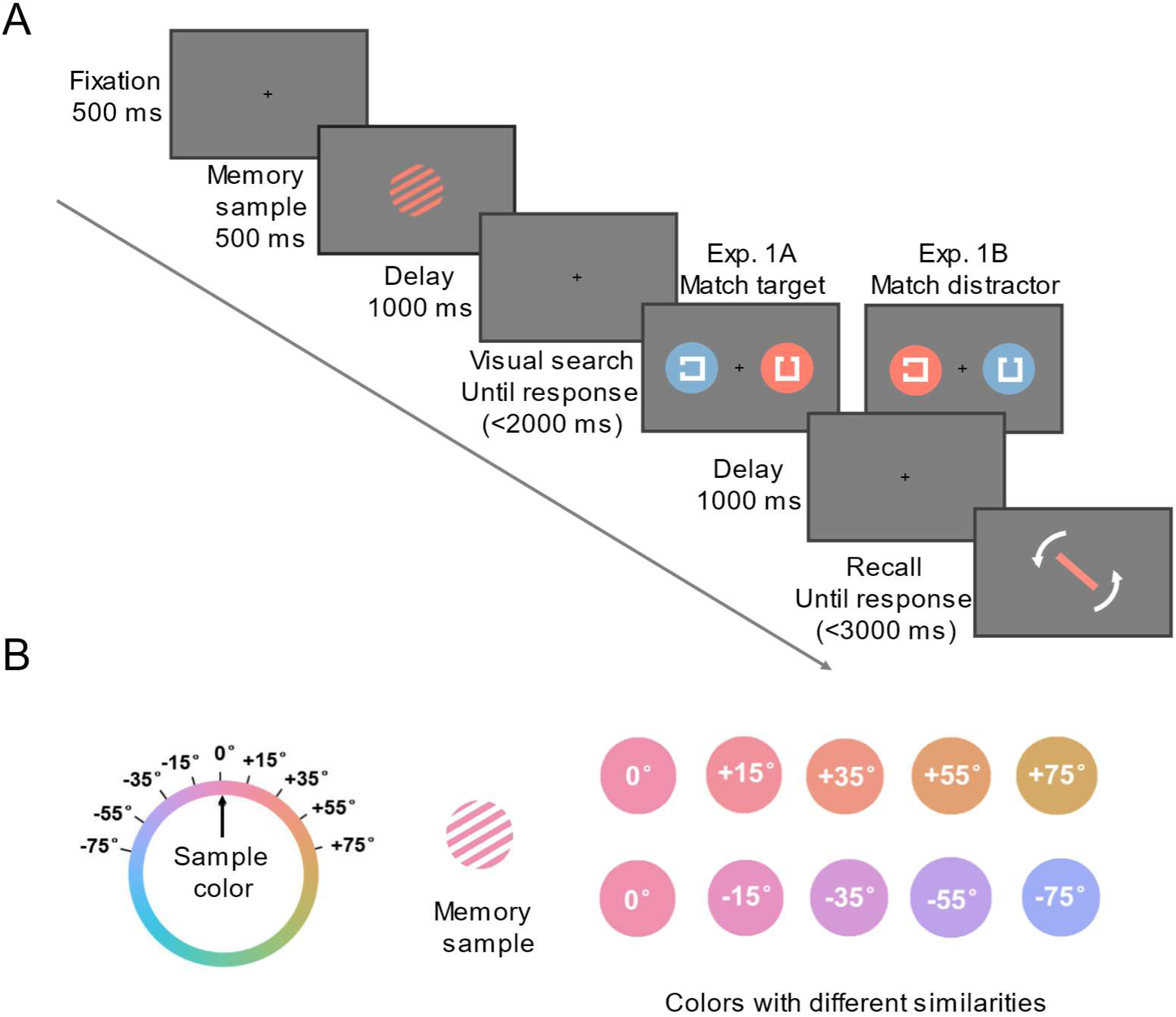
Procedure and example stimuli for Experiment 1. **A:** Examples of experimental trials in Experiment 1. While remembering the orientation of the memory sample, the participants were instructed to indicate whether the gap of the target was up or down during the delay period. Importantly, the color of the background behind the search target (Experiment 1A) or the distractor (Experiment 1B) matched the color of the memorized grating (i.e., VWM-irrelevant feature) with five levels of feature similarity. **B:** Example of a memory sample and the corresponding background color in the Landolt-C search task with different feature similarities (0°, 15°, 35°, 55°, or 75°) from the color of the memory sample in the CIE L*a*b color space.

### Design

The feature similarity between the color of the memory sample and the background color of the search target (Experiment 1A) or distractor (Experiment 1B) was manipulated at five varying levels. The feature similarity was defined by the angle degree in the CIE L*a*b* color space, as illustrated in Fig. 1B (5 similarities: 0°, 15°, 35°, 55°, or 75° with clockwise or counterclockwise rotation directions related to the color of the memorized sample; a larger angle represents lower similarity). To mitigate the potential interaction between the target and distractor, the background colors of the visual search targets and distractors were consistently set 180° apart on the color wheel. In both Experiments 1A and 1B, each participant completed 10-30 practice trials and ten blocks of 400 experimental trials. The proportion of each of the five similarity conditions was the same, resulting in 80 trials per similarity condition. All trial types were randomly interleaved.

### Data analysis

#### Search performance

Trials with no response, incorrect responses, or response times larger than 3 standard deviations from the mean were excluded from the dataset. In addition, the distribution of recall errors (angular difference between response and memory samples) was also considered. Any trial with a recall error that was outside 3 standard deviations of the recall error distribution of all trials was categorized as a forgotten trial and subsequently eliminated. This resulted in a data loss of 2.9% in Experiment 1A and 2.3% in Experiment 1B. The remaining trials were included in further analyses.

We conducted two types of analyses to assess the modulation pattern of similarity on the WM- captured attention using JASP (version 0.16.3) and MATLAB. In the first analysis, we calculated the search reaction time for each similarity level by averaging the clockwise (e.g., +15°) and counterclockwise (e.g., -15°) deflected conditions. We subsequently conducted a one-way repeated- measures ANOVA (5 similarity levels: 0°, 15°, 35°, 55°, and 75° for the color feature) and multiple pairwise comparisons (paired-samples two-tailed t tests). The *p* values of multiple pairwise comparisons were corrected via Benjamini and Hochberg’s FDR-controlling method (Benjamini & Hochberg, 1995).

In the second analysis, we aimed to determine whether a monotonic or nonmonotonic model provided a better fit for the attentional capture effect. This allowed us to further quantify the modulation pattern of similarity. First, we calculated the attentional capture effect as the normalized difference between each similarity level and the baseline condition (the most dissimilar condition).

In Experiment 1A, Normalized 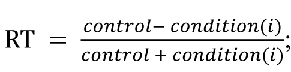 in Experiment 1B, Normalized RT =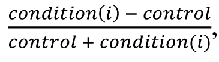 where 𝑖 represents each similarity condition, with values ranging from 1 to 7 (-55°, -35°, -15°, 0°, 15°, 35°, 55°). To capture the full profile, we calculate the normalized RT separately for each clockwise-deflected and counterclockwise-deflected similarity level. The monotonic model was implemented as a Gaussian function to examine whether the modulation profile was consistent with the neuronal tuning curve profile in the early visual cortex. There are three free parameters in the Gaussian function:

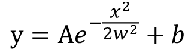

where y is the normalized search response time; 𝑥 is the similarity between the irrelevant feature in VWM and the search background; and the shape of the function is determined by the free parameters A, 𝑤, and 𝑏 . The non-monotonic model was implemented as a difference of Gaussian (DoG) function. The DoG function has five free parameters:

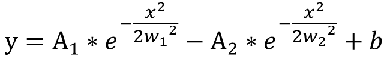

where A_l_, A_2_, 𝑤_l_, 𝑤_2_ and 𝑏 are the five free parameters that determine the shape of the function. To assess the evidence in favor of each model, we refer to the methods utilized in a previous study (Fang et al., 2019; Fang & Liu, 2019). We calculate both the Bayesian information criterion (BIC; (Schwarz, 1978) and the Akaike information criterion (AIC; (Akaike, 1998), assuming a normal error distribution. The BIC is defined as follows:

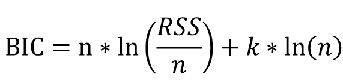

where n is the number of observations, 𝑘 is the number of free parameters, and 𝑅𝑆𝑆 is the residual sum of squares. A smaller BIC indicates a better fit. Additionally, we calculate the Bayes factor (BF) of the Gaussian function over the difference of the Gaussian function on the basis of the BIC approximation (Wagenmakers, 2007):

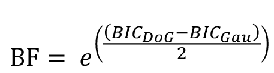

where 𝐵𝐼𝐶_DoG_ represents the difference in the Gaussian function fitting results and where 𝐵𝐼𝐶_Gau_

represents the Gaussian function fitting results.

Apart from the Bayesian information criterion (BIC), we also calculate the Akaike information criterion (AIC), which penalizes model complexity more slightly. The AIC is defined as follows:

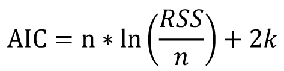

where n is the number of observations, 𝑘 is the number of free parameters, and 𝑅𝑆𝑆 is the residual sum of squares. To compare the models, we calculated the likelihood ratio of the Gaussian function over the difference of Gaussian function (Burnham & Anderson, 2002).

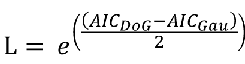

where 𝐴𝐼𝐶_DoG_ represents the difference in the Gaussian function fitting results and where 𝐴𝐼𝐶_Gau_ represents the Gaussian function fitting results.

#### Memory performance

To quantify the performance of the memory task, we employed the standard mixture model (Zhang & Luck, 2008) to fit our data, which decomposes the recall error distribution into a mixture of two distributions: a von Mises distribution (circular normal distribution) and a uniform distribution. The von Mises distribution is defined by two parameters: the distribution’s mean, which represents the overall recall shift from the correct feature value, and the standard deviation (𝑆𝐷) of the distribution, which represents memory precision (a larger SD corresponds to lower memory precision). When memory features are still retained in the VWM, responses may follow a von Mises distribution. However, subjects might forget the feature and report with guesses so that each feature value has an equal probability of being chosen, resulting in a uniform distribution. The uniform distribution is characterized by a single parameter, 𝑔, which represents the height of the uniform distribution and signifies the guess rate. The mean, standard deviation and guess rate were calculated via Memtoolbox (Suchow et al., 2013) in MATLAB. One-way repeated-measures ANOVA was subsequently performed on each of these three indices to examine the influence of feature similarity on working memory performance.

Some researchers have highlighted that the statistical power of employing a standard mixture model to detect an inherent change in precision depends on the baseline memory performance level. To be more specific, statistical power is maximized when precision is high and the guess rate is low (Rademaker et al., 2018). In the present study, participants were tasked with memorizing only one item per trial, thereby maximizing the baseline memory performance.

### Transparency and openness

All data, analysis and task codes have been made publicly available via the Open Science Framework at https://osf.io/3vxqt/. Dada were analyzed using MATLAB and JASP version 0.16.3. This study was not pre-registered.

## Results

### Search performance

Figure 2 depicts the search reaction times (RTs) in Experiment 1. In Experiment 1A (Fig. 2A), one-way repeated measures analysis of variance (ANOVA) with factor similarity (five levels) was conducted, revealing a significant main effect of similarity (*F* (4,108) = 12.47, *p* < 0.01, 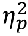 = 0.32). Multiple pairwise comparisons, corrected via Benjamini and Hochberg’s FDR-controlling method (Benjamini & Hochberg, 1995), revealed significant differences (FDR-corrected threshold = 0.04) between 0° and 35° (*t* (27) = -3.25, *p* < 0.01, Cohen’s *d* = -0.61), 0° and 55° (*t* (27) = -3.25, *p* < 0.01, Cohen’s *d* = -0.61), 0° and 75° (*t* (27) = -5.18, *p* < 0.01, Cohen’s *d* = -0.98, 15° and 35° (*t* (27) = -2.61, *p* < 0.05, Cohen’s *d* = -0.49), 15° and 55° (*t* (27) = -3.06, *p* < 0.01, Cohen’s *d* = -0.58), 15° and 75° (*t* (27) = -5.28, *p* < 0.01, Cohen’s *d* = -1.00), 35° and 75° (*t* (27) = -2.92, *p* < 0.01, Cohen There were no significant differences between 0° and 15° (*t* (27) = -0.85, *p* = 0.40, Cohen’s *d* = -0.16) or between 35° and 55° (*t* (27) = -0.83, *p* = 0.42, Cohen’s *d* = -0.16). These results showed that the search speed increased when the color of the target background matched the irrelevant feature in VWM, and the degree of VWM-biased attention decreased monotonically as feature similarity decreased.

**Fig. 2.**
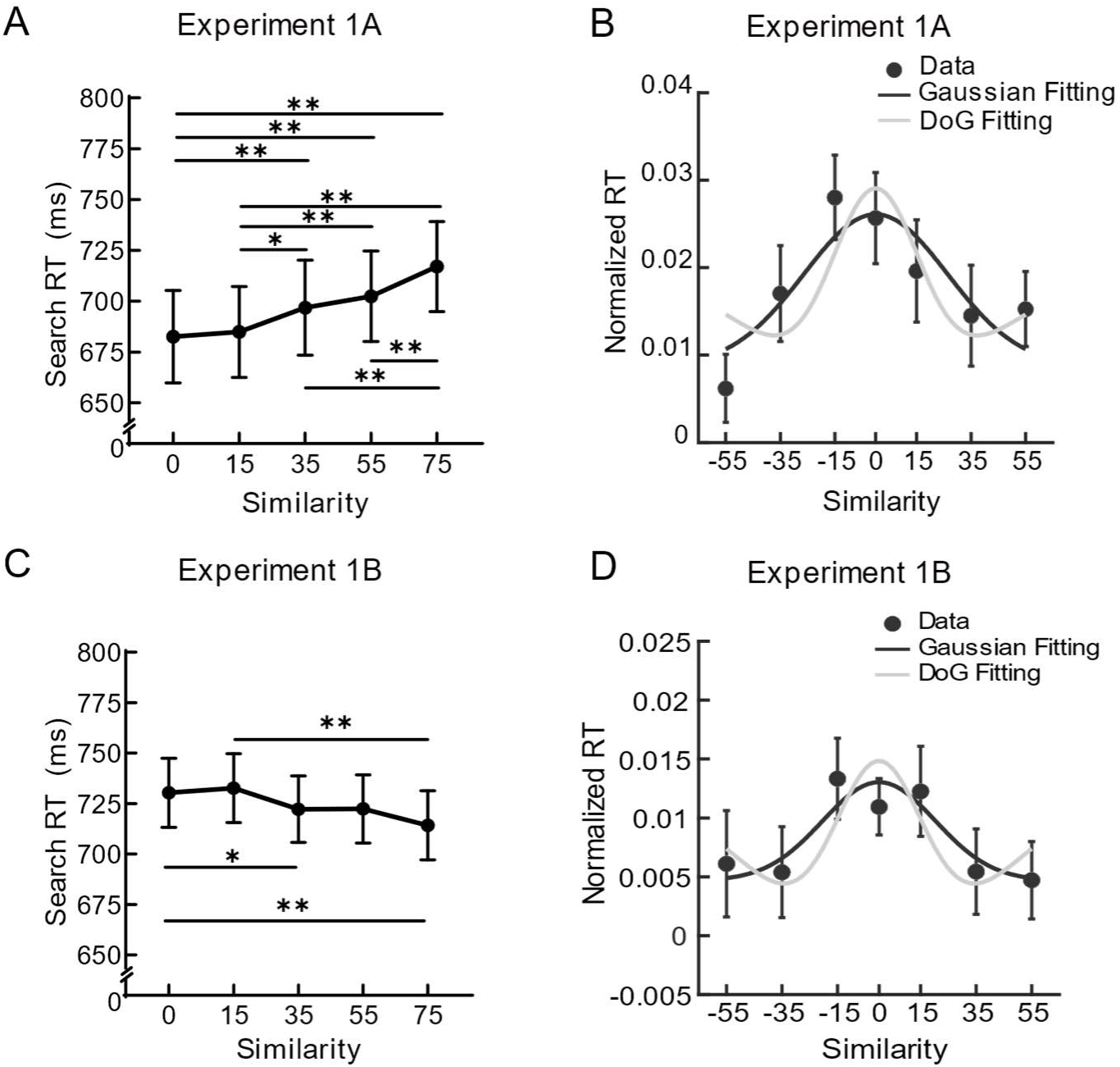
Results for Experiment 1. Panels (**A**) and (**C**) show the search reaction times (RTs) when the backgrounds of the search target (**A**) and distractor (**C**) were identical or similar to the memory-irrelevant feature in Experiments 1A and 1B, respectively. RTs for each level of similarity were averaged across the clockwise (e.g., +15°) and counterclockwise (e.g., -15°) directions. The search speed increased when the target matched the irrelevant feature and decreased when the distractor matched it. As the feature similarity decreased, the degree of VWM-biased attention decreased monotonically in both Experiments 1A and 1B. Panels (**B**) and (**D**) show the normalized RTs as a function of feature similarity (filled symbols) and the model-fitting results using both the monotonic Gaussian function (black line) and the non-monotonic difference of Gaussian function (gray line) in Experiments 1A and 1B, respectively. Note that normalized RTs were calculated by subtracting the least dissimilar condition (i.e., 75°) from each similarity level in Experiment 1A, whereas was calculated by subtracting each similarity level from the least dissimilar condition in Experiment 1B. Data points represent group means. The error bars represent the standard errors of the means (SEMs). \**p* &lt; 0.05, \*\**p* &lt; 0.01, uncorrected.

To quantify the overall profile of this VWM-biased attention, we fitted a monotonic model (i.e., Gaussian function) and a non-monotonic model (i.e., difference of Gaussian function, DoG) to the group-averaged normalized reaction time (Fig. 2B). Within the -55°--55° offset range, the Gaussian function (R^2^ = 0.75, BIC = -73.79) exhibited a stronger preference than did the DoG function (R^2^ = 0.58, BIC = -65.30), with a Bayes factor of 70.09. The AIC evidence also revealed that the Gaussian function (AIC = -73.63) was 73.99 times more likely than the DoG function (AIC = -65.02). This result provides evidence that the feature profile of WM-biased attention was a monotonic gradient profile resembling the tuning curve of feature-selective neurons in the early visual cortex.

In Experiment 1B, one-way repeated-measures ANOVA revealed a significant main effect of similarity, *F* (4,104) = 4.17, *p* < 0.01, 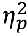= 0.14, which suggests that memory-search similarity modulates VWM-biased attention even when it matches the distractor. Multiple pairwise comparisons (FDR-corrected threshold = 0.02) revealed significant differences between 0° and 35° (*t* (27) = 2.57, *p* < 0.02, Cohen’s *d* = 0.49), 0° and 75° (*t* (27) = 4.27, *p* < 0.01, Cohen’s *d* = 0.82), 15° and 75° (*t* (27) = 3.78, *p* < 0.01, Cohen’s *d* = 0.73). The other pairwise comparisons were not significant (*ps* > 0.07; more detailed statistics can be found in Supplementary Table 1, Additional file 1). More importantly, we fitted both the Gaussian and DoG models to the group-averaged normalized RTs as in Experiment 1A (Fig. 2D). The Bayes factor (260.70) strongly supported the Gaussian function (R^2^ = 0.81, BIC = -85.41), surpassing the difference of the Gaussian function (R^2^ = 0.48, BIC= -74.29). Similarly, the likelihood ratio (275.19) based on the AIC also supported the Gaussian function (AIC = -85.26) rather than the difference Gaussian function (AIC = -74.02). These results indicated that irrelevant features in VWM could interfere with attention allocation to incoming information even when they are always matched with search distractors, and the degree of this VWM-biased attention decreased as the feature similarity between the color of the distractor background and the color of the memorized grating decreased.

To rule out the alternative possibility that VWM-biased attention to reaction time reflects the speed‒accuracy trade-off, we performed an ANOVA test on accuracy. The accuracy remained consistently high across all similarity levels for both Experiments 1A and 1B (all > 99.10%). The results revealed no significant main effect of similarity (Experiment 1A: *F* (4,108) = 0.97, *p* = 0.43, 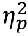 = 0.04; Experiment 1B: *F* (4,104) = 1.20, *p* = 0.32, 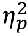 = 0.04), suggesting that the difficulty of p p the search task was not affected by similarity and eliminating the speed‒accuracy trade-off.

### Memory performance

To quantify the performance of the memory task, we focused on two critical indices for each experiment, the mean and standard deviation of the von Mises distribution, which reflect the precision and bias of the memory from the actual stimuli. One-way repeated measures ANOVAs with the factor similarity conducted separately for each parameter of the recall performance showed no significant main effect of similarity for all parameters in both Experiments 1A and 1B (all *ps* > 0.31). These results showed that participants exhibited consistent performance across all similarity levels, which is to be expected given that the interim search task involves only the memory- irrelevant color but not the to-be-remembered orientation.

## Discussion

Although the context of the search display was completely irrelevant to the search task, the search speed was fastest when the target background matched the memory-irrelevant feature and slowest when the distractor matched the irrelevant feature. This observed VWM-biased attention effect is consistent with previous findings and suggests that irrelevant features can be encoded in VWM (Foerster, 2017; Gao et al., 2016; Soto & Humphreys, 2009). More importantly, we further revealed that the degree of VWM-biased attention decreases monotonically with decreasing similarity. The profile of the VWM-biased attention followed a monotonic function, which is in line with the tuning properties of neurons in the early visual cortices. These results imply that irrelevant features are retained in sensory cortices and represented as sensory analogs.

## Experiment 2: VWM-biased perception in the discrimination task

In Experiment 1, the sensory analog representation of VWM was inferred from the profile VWM-biased attention. However, one might argue that the monotonic gradient profile observed in Experiment 1 may not be a result of working memory’s modulation of perceived information but rather a result of working memory’s modulation of perception via its interaction with attention. To further examine whether VWM directly influences perception itself, Experiment 2 deployed a dual- task paradigm with an interleaved perceptual discrimination task to capture the influence of irrelevant features on perceptual discrimination.

### Method

#### Participants

Thirty-one (23 females and 8 males; age: M = 18.6 years, SD = 1.6) and thirty-three (24 females and 9 males; age: M = 18.9 years, SD = 2.0) individuals were participated in Experiments 2A and 2B, respectively. However, 4 participants in Experiment 2A and 5 participants in Experiment 2B were excluded because their data did not fit the psychometric function, leaving 27 participants and 28 participants for the two experiments. They provided written informed consent and all reported normal or corrected-to-normal vision with no history of mental illness. They were paid for their participation.

#### Apparatus and stimuli

The memory sample was a colored and oriented grating (2.0° in radius; spatial frequency: 2

cycles/degree). The color of the grating was chosen at random from a set of 20 colors that were sampled at equal intervals (18°/steps over the entire 360° color wheel) from the CIE L*a*b color space. Orientation was chosen from 20 possible orientations, equally spaced at 5° intervals, ranging from 20° to 65° and 115° to 160° to avoid specific orientations such as horizontal and vertical orientations. For the memory report, a colored bar and a color wheel (6° in radius and 0.8° in width) appeared at the center of the display in Experiments 2A and 2B, respectively. For the discrimination stimuli, two gratings (2.0° in radius) were presented simultaneously at an eccentricity of 3.5°. We manipulated the distance between the two discriminative gratings in color (Experiment 2A) or orientation (Experiment 2B) space from 0° to 18° in 3° increments (0°, 3°, 6°, 9°, 12°, 15°, 18°) to construct psychometric functions of color or orientation discrimination tasks. Importantly, under similar conditions, the color/orientation of the two discrimination gratings were always on either side of the VWM irrelevant feature with the same distance; under dissimilar conditions, the two discriminated features were always on either side of the farthest distance (i.e., 180° apart in the color space or 90° apart in the orientation space) relative to the VWM irrelevant feature with the same distance (see Fig. 3B). In the other feature dimension, the perceptually irrelevant feature (i.e., orientation in Experiment 2A or color in Experiment 2B) was kept the same for both discrimination gratings and was systematically manipulated with five levels of similarity (similarity: 0°, 15°, 35°, 55°, 75°) relative to the memorized features.

**Fig. 3.**
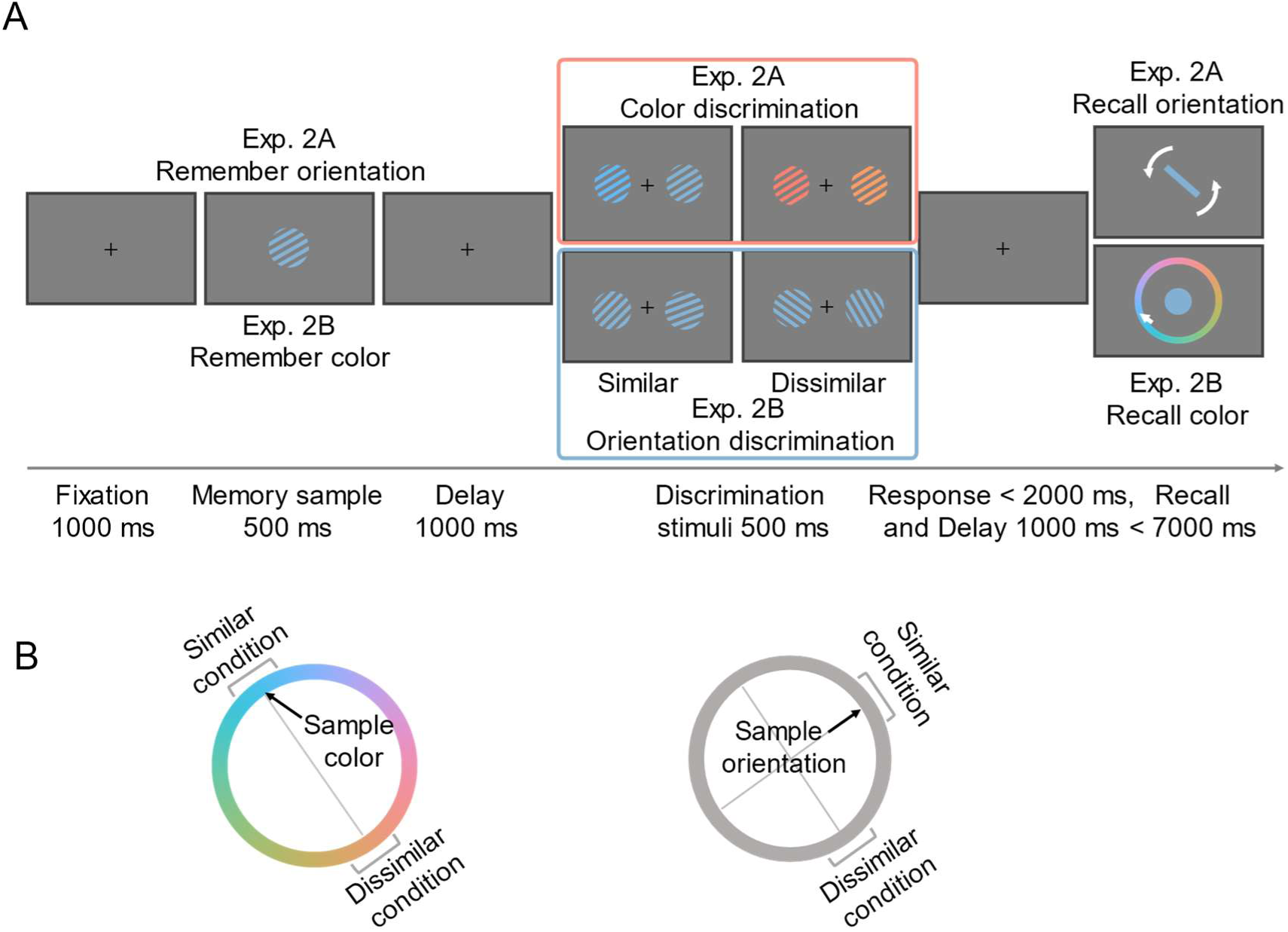
Procedure and example stimuli for Experiment 2. **A:** Procedure in Experiment 2. Participants were required to discriminate the color (Experiment 2A) or orientation (Experiment 2B) of the two perceived gratings during the retention, while memorizing the orientation (Experiment 2A) or color (Experiment 2B) of the memory sample for later recall. That is to say, the memory-irrelevant feature was color in Experiment 2A and orientation in Experiment 2B. **B:** Illustration of the similar and dissimilar conditions in color space and orientation space in Experiment 2. In the similar condition, the discriminative features of the two perceived gratings were equally spaced on either side of the memory-irrelevant feature at varying distances to assess the discrimination thresholds. In the dissimilar condition, they were placed on either side of the largest distance from the feature held in working memory (i.e., 180° apart in the color space or 90° apart in the orientation space) with varying distances. The dissimilar condition served as the baseline condition for assessing the unbiased discrimination thresholds.

#### Procedure

As shown in Fig. 3A, in Experiment 2, each trial began with a 1000 ms fixation display, followed by the sample grating for 500 ms and then a 1000 ms blank delay. The discrimination stimuli were then presented for 500 ms, and the participants had 2000 ms to respond, during which a fixation cross was displayed on the screen. After the response, there was a 1000 ms blank delay, followed by the recall display, which disappeared either after a response or after 7000 ms. Participants were asked to remember the orientation (Exp. 2A) or color (Exp. 2B) of the memory sample and to discriminate whether the color (Exp. 2A) or orientation (Exp. 2B) of two stimuli was the same or not in the interleaved perceptual task during retention. For memory reports, participants in Experiment 2A were required to rotate the vertical bar clockwise or counterclockwise to report the remembered orientation as accurately as possible. In Experiment 2B, participants were asked to respond as accurately as possible by moving the mouse on the color wheel and clicking to confirm the selection. The color wheel was randomly rotated on each trial to prevent participants from converting color memory into spatial location memory. It was stipulated to the participants that they should refrain from using any strategies in the completion of the memory task. Each participant completed 10--30 practice trials and 280 experimental trials. There were 140 trials for both similar and dissimilar conditions. All trials were counterbalanced across the five similarity levels between perceptually irrelevant features and WM-relevant features. All trial types were randomly interleaved. No feedback was provided in either the memory or the discrimination task.

#### Data analysis

***Threshold Fitting:*** The color and orientation discrimination thresholds were estimated by fitting individual data from different conditions to the Weibull function through the Palamedes toolbox (Kingdom & Prins, 2010; Prins & Kingdom, 2018) in MATLAB:

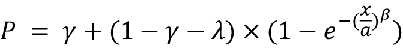

in which 𝑃 represents the proportion of “different” responses in the discrimination task at varying feature value differences 𝑥; 𝛼 and 𝛽 are free parameters representing the threshold and slope of the fit, respectively. 𝛾 is the chance-level performance or guess rate, which indicates the probability of choosing the correct response if an observer was to guess in a trial. The parameter λ is commonly referred to as the lapse rate. This rate indicates that the observer may respond independently of the stimulus level on a small proportion of trials, possibly owing to a momentary lapse of attention or a sneeze. Both 𝛾 and λ were also set as free parameters. We chose the point at which the subjects were 75% likely to report “different” as their discrimination threshold. To ascertain whether discrimination thresholds differed between different conditions, paired-samples t tests (two-tailed) were employed in Experiment 2.

***Memory performance***: In Experiment 2, the analysis of memory task performance was identical to that in Experiment 1. In Experiment 3, we conducted two-way repeated measures ANOVAs (delay × similarity) on the same parameters.

## Results

### Discrimination threshold in the perceptual task

The participants were asked to remember the orientation (Experiment 2A) or color (Experiment 2B) of a multifeature object while discriminating the memory-irrelevant feature (i.e., color in Experiment 2A and orientation in Experiment 2B) of two gratings during the delay period of the VWM task. Experiment 2B was conducted to characterize VWM-biased perception for the orientation feature dimension to determine the generality of our findings. The feature similarity between the memory-irrelevant feature and the discriminative feature was manipulated (Fig. 3A). The two discriminative gratings were equally spaced on either side of the memory-irrelevant feature and the most dissimilar feature (180° in color space or 90° in orientation space) in similar conditions and dissimilar conditions, respectively (Fig. 3B). We reasoned that if sensory areas are recruited to represent memory-irrelevant features in WM, the neural population of the memory-irrelevant information should overlap with the current visual information when they are highly similar in feature space, making the two discrimination gratings be perceived toward the memorized stimuli and resulting in a higher discrimination threshold related to the dissimilar condition.

To calculate the discrimination thresholds, we fitted the Weibull function for each participant and for each similarity condition separately. As shown in Fig. 4, the perceptual threshold was significantly larger for the similar condition than for the dissimilar condition in both Experiments 2A with the color dimension (*t* (26) = 2.54, *p* < 0.05, Cohen’s *d* = 0.49) and 2B with the orientation dimension (*t* (27) = 4.20, *p* < 0.01, Cohen’s *d* = 0.79), as revealed by paired t tests (two-tailed). This result indicates that the perceived color and orientation were biased toward the memory-irrelevant feature in VWM, which led to an increased threshold in the similar condition compared with the dissimilar condition.

**Fig. 4.**
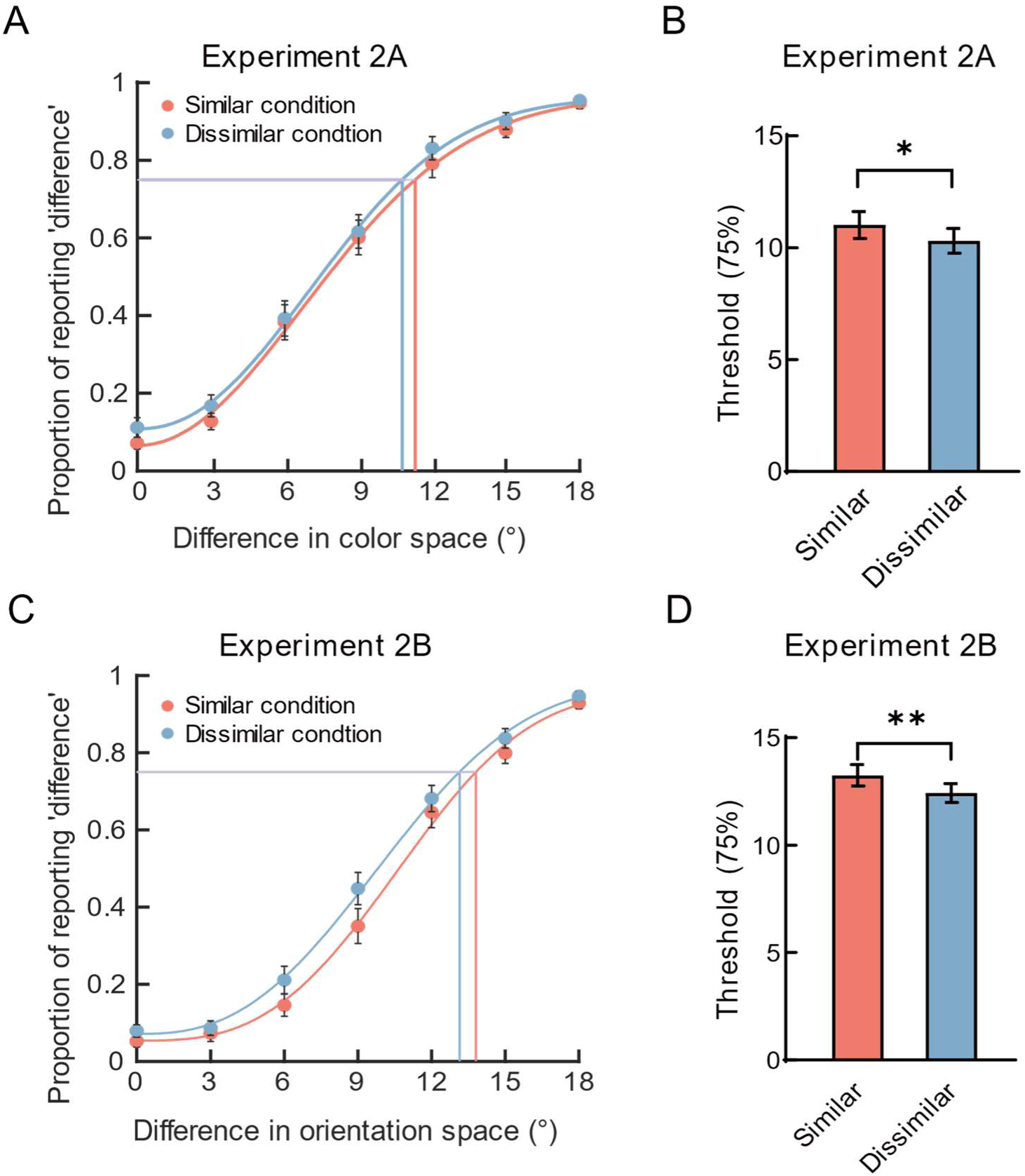
Results of perceptual discrimination in Experiment 2. A: The fitting results of similar conditions and dissimilar conditions in Experiment 2A with the color discrimination task. The filled points represent the average proportion of participants reporting ’differences’ in the perceptual discrimination task for seven color difference levels between the two perceived gratings in color space (0°, 3°, 6°, 9°, 12°, 15°, 18°). The orange and blue lines are the fitted Weibull functions for similar and dissimilar conditions. B: Seventy-five percent color discrimination thresholds plotted by different similarity levels in Experiment 2A. C: The fitting results of similar conditions and dissimilar conditions in Experiment 2B with the orientation discrimination task. D: 75% orientation discrimination thresholds plotted by different similarity levels in Experiment 2B. In panels B and D, the bars represent the group means. The error bars represent the SEMs. \**p* &lt; 0.05, \*\**p* &lt; 0.01.

#### Memory performance in the VWM task

Previous studies have revealed the influence of perceived visual stimuli on the recall performance of memory-relevant features, providing a shared neural representation for ongoing perception and maintaining VWM content(Rademaker et al., 2019; Teng & Kravitz, 2019). Given that the feature dimension of the memory-relevant and perception-relevant feature is orthogonal in the same trial, we could ask whether and how the perceived irrelevant feature biases VWM performance. The degree of similarity between perceptually irrelevant features and VWM-relevant features was manipulated at five different levels. In Experiment 2A, orientation recall performance did not have a significant main effect on either index, as revealed by the repeated measures ANOVAs (all *ps* > 0.09). In Experiment 2B, the SD, but not the mean, increased as the similarity level decreased (*F* (4,108) = 20.84, *p* < 0.01, 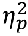= 0.44; see Supplementary Table 5 for pairwise comparisons), suggesting that the perceptually irrelevant feature modulated the memory precision of the feature to be remembered. We replicated previous findings that the degree of modulation increased monotonically with decreasing feature similarity, providing evidence for the sensory representation of the memory-relevant feature in VWM (Teng & Kravitz, 2019).

## Discussion

Using interleaved perceptual discrimination tasks during VWM maintenance, we showed that the perceived stimuli were biased toward memory-irrelevant features, resulting in elevated perceptual thresholds for both orientation and color discrimination features. The VWM-biased perception effect observed in Experiment 2 suggested that the sensory analog representation for memory- irrelevant features was retained in VWM and extended the sensory recruitment theory to memory- irrelevant features. However, some neuroimaging studies have shown no evidence of neural representations of irrelevant features in VWM (Serences et al., 2009; Yu & Shim, 2017). The discrepancy between the results of our study and those of these neuroimaging studies might be due to the length of the delay period (Olivers et al., 2006; Xu, 2010). The memory-irrelevant feature might have decayed (Logie et al., 2011) or might have been coded abstractly (Bae & Luck, 2019) over time. In light of this discrepancy and the critical role of the delay period in understanding the mechanisms of VWM, Experiment 3 was designed to investigate whether the representation of VWM changes over time.

## Experiment 3: The time course of the VWM- perception interaction

Specifically, Experiment 3A examined whether memory-irrelevant feature-biased perception at 1 s and 3 s was delayed after the memory sample disappeared, whereas Experiment 3B examined the same question for the memory-relevant feature. Experiment 3B served as a control experiment to distinguish whether the lack of VWM-biased perception at longer delays occurred for the entire object or only for the irrelevant feature. The participants were asked to maintain the color of the sample grating for later recall while performing an orientation discrimination task during the delay period in Experiment 3A while maintaining the orientation while performing the same orientation discrimination task in Experiment 3B. Orientation was used as the discriminative feature dimension in Experiment 3 given the larger effect size of VWM-biased perception for the orientation in Experiment 2. The recall tasks were set differently to keep the perceptual discrimination task consistent for irrelevant and relevant features in experiments 3A and 3B, respectively.

### Method

#### Participants

Experiments 3A and 3B included thirty-one (22 females and 9 males; age: M = 19.8 years, SD = 0.9) and thirty (25 females and 5 males; age: M = 19.8 years, SD = 1.3) participants, respectively. As the data did not align with the psychometric function, 5 participants were excluded from the analysis in Experiments 3A and 3B. This resulted in final sample sizes of 26 and 25 participants in Experiments 3A and 3B, respectively. They provided written informed consent and all reported normal or corrected-to-normal vision.

#### Apparatus, Stimuli and Procedure

The stimuli employed in Experiment 3A were identical to those utilized in Experiment 2B, with the exception that the seven orientation difference levels of the two discrimination gratings were 0°, 6°, 9°, 12°, 15°, 18° and 21°. The stimuli used in Experiment 3B were similar to those used in Experiment 3A. The only difference between these two experiments was the recall task with the color recall task and orientation recall task in Experiments 3A and 3B, respectively.

The procedure of Experiment 3A was similar to that of Experiment 2B, with the exception that the delay following the presentation of the memory sample was 1000 ms or 3000 ms and the delay after discrimination was 500 ms. The sole distinction between the procedures of Experiments 3B and 3A pertains to the memory task, whereby in Experiment 3B, participants were required to recall the orientation of the sample grating, but not its color. In both Experiments 3A and 3B, each participant completed 560 experimental trials.

#### Data analysis

***Perceptual threshold***: The data analysis in Experiment 3 was the same as Experiment 2. Two-way repeated measures ANOVAs with delay and similarity were conducted for perceptual discrimination task.

***Memory performance***: The analysis of memory task performance was identical to that in experiments 1 and 2. In Experiment 3, we conducted two-way repeated measures ANOVAs with delay and similarity on the same parameters.

## Results

### Discrimination threshold in the perceptual task

Perceptual discrimination thresholds were calculated by fitting psychometric functions for each condition and each experiment (Fig. 5). In Experiment 3A, two-way (2 delays × 2 similarities) repeated-measures ANOVA on perceptual thresholds revealed significant main effects of delay (*F* (1,25) = 5.49, *p* < 0.05, 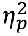 = 0.18) and similarity (*F* (1,25) = 13.70, *p* < 0.01, 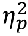 = 0.35), as well as a significant interaction effect between these factors (*F* (1,25) = 4.15, *p* = 0.05, 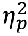 = 0.14). Further simple effect analysis revealed that the discrimination threshold was significantly greater for the similar condition than for the dissimilar condition at a 1000 ms delay (*t* (25) = 4.39, *p* < 0.01, Cohen’s *d* = 0.86; Fig. 5B) but not at a 3000 ms delay (*t* (25) = 1.54, *p* = 0.14, Cohen’s *d* = 0.30). These results indicated that, for the irrelevant feature, VWM-biased perception disappeared in the prolonged condition.

**Fig. 5.**
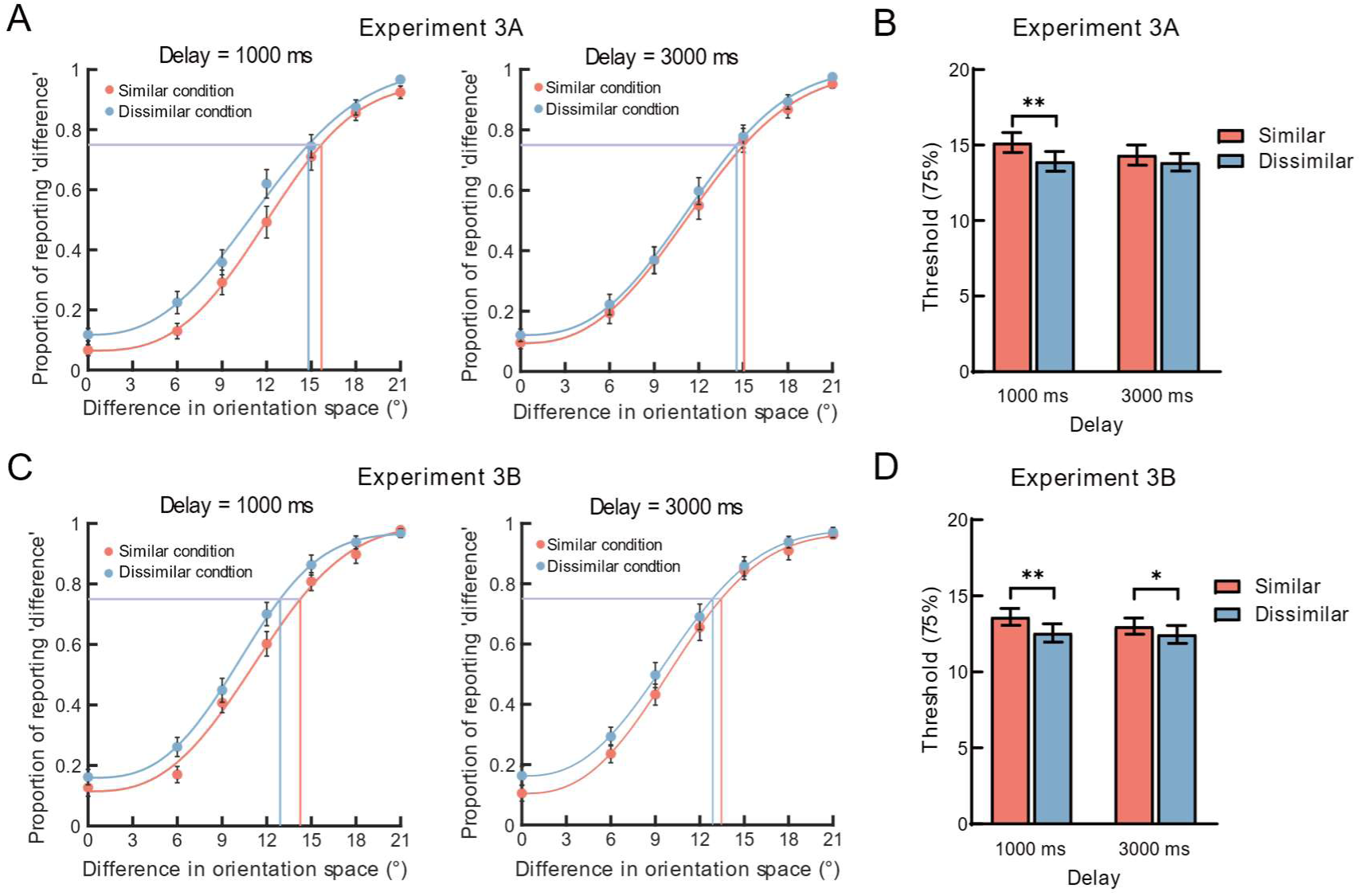
Results of the orientation discrimination task in Experiment 3. **A:** The fitting results for similar and dissimilar conditions, plotted separately for different delays in Experiment 3A with orientation discrimination and color memory. **B:** The orientation discrimination thresholds plotted by delay and similarity in Experiment 3A. **C:** The fitting results for similar conditions and dissimilar conditions, plotted separately for different delays in Experiment 3B when the discrimination and memory dimensions were both oriented. **D:** The orientation discrimination thresholds plotted by delay and similarity in Experiment 3B. The data points in panels A and C represent the average proportion of reported ‘differences’ across all participants in the perceptual discrimination task for seven orientation difference levels. The solid lines in panels A and C show the fitted Weibull functions used to determine the discrimination threshold. In panels B and D, the bars represent the group means. \**p* &lt; 0.05, \*\**p* &lt; 0.01. The error bars represent the SEM.

In Experiment 3B, two-way (2 delays × 2 similarities) repeated-measures ANOVA revealed a significant main effect of similarity (*F* (1,25) = 13.81, *p* < 0.01, 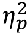 = 0.37), with a higher discrimination threshold for the similar condition than for the dissimilar condition, whereas there was neither a significant main effect of delay (*F* (1,25) = 3.21, *p* = 0.08, 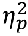 = 0.12) nor an interaction effect between delay and similarity (*F* (1,25) = 1.49, *p* = 0.24, 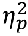= 0.06). Further paired t tests confirmed that the threshold of the similarity condition was greater than that of the dissimilar condition, both for 1000 ms (*t* (25) = 3.20, *p* < 0.01, Cohen’s *d* = 0.64) and 3000 ms (*t* (25) = 2.03, *p* = 0.05, Cohen’s *d* = 0.41) delays (Fig. 5D). The results implied that VWM-biased perception occurred at both short and long delays.

#### Memory performance in the VWM task

In Experiment 3A, we also tested whether the modulation of the perceptually irrelevant feature on the VWM differed for different similarity levels and different delay periods. We performed two two-way (2 delays × 5 similarities) repeated measures ANOVAs on the SDs and means separately. We found that the SD was smaller for the 1 s delay period than for the 3 s delay period, as indicated by the significant main effects of delay (*F* (1,25) = 11.54, *p* < 0.01, 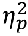 = 0.32). The main effect of similarity was also significant (*F* (4,100) = 41.38, *p* < 0.01, 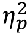 = 0.62), but there was no significant interaction effect between the two (*p* = 0.20). Post hoc analysis revealed that the SD increased monotonically as the similarity between the perceived irrelevant feature and the memory-relevant feature decreased (detailed statistics can be found in Supplementary Table 5). For the mean, both the main effects and the interaction effect were not significantly different (*p* > 0.10). These results indicate that memory precision was enhanced when the color of the memory sample was highly similar to the perceived color, even when the color was irrelevant to the perceptual discrimination task. In addition, the memory precision decreases with increasing delay time, which also replicates the previous finding that memory performance decreases with time (Rademaker et al., 2018).

In Experiment 3B, we also conducted two-way (2 delays × 2 similarities) repeated-measures ANOVAs for each index. The results revealed that the SD for the similar condition was significantly smaller than that for the dissimilar condition (*F* (1, 24) = 12.72, *p* < 0.01, 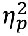 = 0.35) and was significantly smaller for the 1000 ms delay condition than for the 3000 ms delay condition (*F* (1,24) = 16.48, *p* < 0.01, 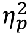 = 0.407), whereas the interaction effect was not significant (*p* = 0.36). The ANOVA of the means revealed that none of the main effects or the interaction effects were significant (all *p*s > 0.10). These results indicate that memory precision was enhanced for similar features and for short delay periods when memory-relevant features and perceptual-relevant features were identical. Moreover, the memory precision decreased as the delay period increased and as the feature similarity between the perceptual- and VWM-relevant features decreased.

## Discussion

The current experiment revealed that memory-irrelevant features biased perception at a 1 s delay but not at a 3 s delay, suggesting the decay of irrelevant information in VWM over time. However, it might be argued that the lack of VWM-biased perception for irrelevant features is caused by the transferred representation in high-level abstract (e.g., verbal) form but not decayed. We rule out this alternative explanation with the presence of VWM-biased perception in VWM at both 1 s and 3 s delays, suggesting the sensory representation of memory-relevant features at both delays. To the best of our knowledge, there is no evidence that an irrelevant feature is transferred to a different code before the relevant feature is transferred. In contrast, there is evidence that the decay of the irrelevant features is faster than that of the relevant feature (Bocincova & Johnson, 2019). Given these clues, our findings suggest that the memory-irrelevant features are represented as sensory analogs for a short period of time.

## General Discussion

The current study investigated the representational nature of VWM by testing how and when memory-irrelevant features alter attentional allocation to and perception of incoming visual information. We found that the irrelevant features in the VWM biased attention regardless of whether they matched the context of the search targets or the distractors. Moreover, this VWM- biased attention decreased monotonically with decreasing similarity between the maintained and perceived task-irrelevant features, which is congruent with the prediction of the sensory representation account. Experiments 2 and 3 tested the second behavioral prediction of the sensory representation account with VWM-biased perception in the discrimination task. We found that the perceived stimuli were biased toward memory-irrelevant features when these two features were highly similar, resulting in a higher perceptual threshold in the similar condition at short delay. Thus, these findings indicate the sensory analog representation of memory-irrelevant features, extending the sensory recruitment theory to memory-irrelevant features.

### Sensory representation of the irrelevant features in visual working memory

Although some studies have reported negative results (Olivers et al., 2006; Sala & Courtney, 2009), accumulating evidence has shown that memory-irrelevant features can bias attention (Foerster, 2017, 2020; Foerster & Schneider, 2019; Gao et al., 2016; Jung et al., 2020; Soto & Humphreys, 2009). Extending these findings, we found that the degree of VWM-biased attention decreased monotonically as the feature similarity between the maintained and perceived stimuli decreased, revealing a monotonic gradient feature profile of VWM-biased attention. Given the origin of the monotonic gradient tuning curve of simple features in the visual cortex, where they are initially processed, our results imply a sensory representation for irrelevant features in the early visual cortex. The sensory analog representation account has been verified with the feature profile or spatial profile for the memory-relevant features in VWM (Fang et al., 2019; Teng & Kravitz, 2019). The present study revealed the feature profiles for the irrelevant features in VWM. In addition, we found VWM- biased perception with the interleaved discrimination task, providing further evidence for sensory analog representation with different perceptual tasks. Using a paradigm similar to ours, recent studies revealed a bidirectional interaction between relevant features in VWM and in perception (Teng & Kravitz, 2019; Teng & Postle, 2021). We provide further evidence that VWM-biased perception differs between memory-irrelevant and memory-relevant features over time. In short, we consistently observed the VWM-biased perception phenomenon in both the interleaved search and discrimination tasks, which is in accordance with the behavioral prediction of the sensory recruitment model of WM (D’Esposito & Postle, 2015; Postle, 2015, 2016). The present study extends sensory recruitment theory to irrelevant features in VWM.

Previous studies have reported mixed results regarding whether irrelevant features of the memorized object are represented in the brain (Bocincova & Johnson, 2019; Serences et al., 2009; Xu, 2010; Yu & Shim, 2017). Some studies revealed the above-chance classification accuracy for irrelevant features at the early delay period (Bocincova & Johnson, 2019; Xu, 2010), whereas other studies reported the absence of VWM-biased attention and above-chance classification for VWM content at prolonged delays (Han & Kim, 2009; Olivers et al., 2006; Serences et al., 2009; Yu & Shim, 2017). The discrepancy between these results might be due to the length of the delay period (Logie et al., 2011) or the task demand of the memory task (Xu, 2010). Consistent with this interpretation, we found that irrelevant feature-biased perception was short-lasting and was not sustained over a longer delay period, suggesting that the irrelevant feature was temporally represented as a sensory analog and subsequently biased ongoing perception. Our results are also consistent with those of other behavioral or eye-tracking studies with various VWM paradigms, which suggested the encoding of task-irrelevant features (Marshall & Bays, 2013; Shen et al., 2013; Shin & Ma, 2016; Yin et al., 2012; Zokaei et al., 2019). Another possibility is that the irrelevant features are represented in an “activity-silent” state via short-term synaptic plasticity without showing persistent delay activity (Masse et al., 2020; Mongillo et al., 2008), which could be difficult to detect with a decoding method. Further neurobiological or neuroimaging studies using advanced techniques are needed to distinguish between these possibilities.

Another related issue is the neural mechanism of VWM. Previous neurophysiological and neuroimaging studies have shown that high-level areas such as frontoparietal cortical areas are critical for VWM (Bettencourt & Xu, 2016; Christophel et al., 2012; D’Esposito & Postle, 2015; Mendoza-Halliday et al., 2024; Sreenivasan & D’Esposito, 2019), whereas others suggest that low- level areas such as visual cortices are necessary for VWM representation (Harrison & Tong, 2009; Huang et al., 2024; Jia et al., 2021; Rademaker et al., 2019; Serences et al., 2009). Here, we also found that relevant features in VWM persistently biased perception throughout the delay period and that the perceived information biased the memorized stimuli, suggesting a sensory representation of relevant features and their subsequent interaction with VWM and perception. Extending these findings, we showed that irrelevant features could interact with the incoming percept, suggesting the involuntary involvement of sensory cortical areas in VWM. Notably, this result does not exclude the involvement of high-level areas in the VWM representation. Fronto-parietal cortical areas have been shown to represent high-level abstract information, such as task rules and categorization(Ester et al., 2015; Lee et al., 2013). It is likely that the fronto-parietal cortical areas play a top-down modulatory role, sending high-level abstract information back to the sensory cortical areas to complete the maintenance of fine information. Indeed, we found that the relevant feature, but not the irrelevant feature, induced VWM-biased perception with a longer delay. It is possible that the feedback signal from the frontoparietal to the sensory area removes the memory-irrelevant information, which requires additional time. Future studies could use on-line TMS (transcranial magnetic stimulation) to examine the temporal order of the critical involvement of frontoparietal and sensory areas in VWM representations for each feature according to task demands.

### The representation of objects vs. features in visual working memory

The results of the current study are relevant to our understanding of the fundamental units of VWM. It has long been unclear whether the fundamental units of VWM are object-based or feature-based. The object-based representation account proposes that multifeature objects are stored as integrated features (Luck & Vogel, 1997; Shin & Ma, 2016), whereas the feature-based representation account proposes that objects are maintained as independent individual features (Bays et al., 2011). When addressing this issue with various testable hypotheses, previous research has reported mixed results (Luria & Vogel, 2011; Shen et al., 2013). These questions have been investigated in multiple aspects, one of which is whether all features, including the memory-irrelevant features, are retained automatically (Bays et al., 2011). Our results shed new light on this issue. With our dual-task paradigm, we can probe the maintenance of irrelevant features with an interim discrimination task. If all features of the object are maintained as integrated rather than separated features, then the irrelevant features would involuntarily influence the incoming perception when the relevant features are doing so, as indicated by a shift in the attention allocation and perceptual threshold of the stimuli that match the irrelevant features. In fact, the present study showed that both the relevant and irrelevant features induced VWM-biased perception at short delays, supporting the object-based account for VWM during the early delay period. At longer delays, we find that only the relevant feature induces VWM-biased perception, suggesting that only the feature that is important for VWM tasks is maintained. These results suggest that both object- and feature-based representations may be true in different situations, explaining previous discrepancies in the literature. The representational unit of VWM may be object-based in the early phase, whereas it may be feature- based in the later phase. Consistent with this speculation, an EEG study reported above chance decoding accuracy for both the memory-relevant and memory-irrelevant features in the early phase but only for memory-relevant features in the later phase (Bocincova & Johnson, 2019). Using N2pc as an electrophysiological index, a previous study revealed that the amplitude is equal between combined objects and separated features at an early stage but a larger amplitude for combined objects than for separated features at a later stage, providing evidence for object- and feature-based attentional selection at different stages (Eimer & Grubert, 2014). Further EEG studies are needed to provide more evidence for the possible dynamics of different representational units.

Another possibility is that all the features are encoded and maintained in VWM but are maintained differently depending on their relevance to the task. Using the surprise test, previous behavioral studies reported that irrelevant features can be recalled but with much lower fidelity than the same feature is the feature to be remembered, suggesting different representations for relevant and irrelevant features (Shin & Ma, 2016; Swan et al., 2016). Previous theoretical accounts of WM have proposed the hierarchical representation hypothesis and the noisy-memory framework. The former hypothesis proposes that the unit of VWM is a hierarchical feature bundle: the top level represents the integrated object, and the bottom level represents individual low-level features (Brady et al., 2011). The noisy-memory framework proposes that the WM representation consists of the packaged resource representing all the features of objects and the target resource representing the relevant features of the memorized content (Shin & Ma, 2017). The different representation modes of relevant and irrelevant features might cause different durations that can be maintained. It will be interesting for future studies to test these hypotheses by systematically comparing the similarities and differences between content and context features in VWM via the same paradigm.

## Conclusions

The current study investigated VWM-biased perceptions of memory-irrelevant features in interleaved visual search and discrimination tasks with concurrent working memory tasks. In conclusion, we showed that VWM-biased attention has a monotonic gradient profile and that VWM content temporally interacts with ongoing visual information, suggesting the sensory analog representation of irrelevant features. These findings are consistent with the predictions of the sensory recruitment model and extend this theory to irrelevant features in VWM. The present study provides behavioral evidence for the representational nature of irrelevant features in VWM and enriches the understanding of the cognitive mechanism of VWM.

## Declarations

### Ethics approval and consent to participate

The study protocols were approved by the Ethics Committee on Human Experimentation of Shaanxi Normal University (Approval Number: 2020-12-006) following the Declaration of Helsinki. Written informed consent was obtained from the participants before the experiments.

### Consent for publication

Not applicable.

### Availability of data and materials

All data, analysis and task codes have been made publicly available via the Open Science Framework at https://osf.io/3vxqt/. Dada were analyzed using MATLAB and JASP version 0.16.3.

This study was not pre-registered.

## Funding

This work was supported by the National Science and Technology Innovation 2030-Major Project 2022ZD0208200, the National Natural Science Foundation of China (No. 31800910) and the Fundamental Research Funds for the Central Universities (No. GK202301002).

### Authors’ contributions

HS: conceptualization, data curation, formal analysis, validation, visualization, writing-original draft, writing-review and editing; QZ: conceptualization, data curation, formal analysis, validation, visualization, writing-original draft, writing-review and editing; JZ: data curation, formal analysis, writing-original draft; YD: data curation, formal analysis; YW: conceptualization, supervision, writing-original draft and writing-review and editing; YL: conceptualization, funding acquisition, supervision, writing-original draft, and writing-review and editing.

## Acknowledgments

Not applicable

